# Quantitative assessment of anti-gravity reflexes to evaluate vestibular dysfunction in rats

**DOI:** 10.1101/590257

**Authors:** Vanessa Martins-Lopes, Anna Bellmunt, Erin A. Greguske, Alberto F. Maroto, Pere Boadas-Vaello, Jordi Llorens

**Affiliations:** Departament de Ciències Fisiològiques, Institut de Neurociènces, Universitat de Barcelona, Feixa Llarga s/n, 08907 L’Hospitalet de Llobregat, Catalunya, Spain; Institut d’Investigació Biomèdica de Bellvitge, IDIBELL, 08907 L’Hospitalet de Llobregat, Catalunya, Spain; Research Group of Clinical Anatomy, Embryology and Neuroscience (NEOMA), Departament de Ciències Mèdiques, Facultat de Medicina, Universitat de Girona, 17003 Girona, Catalunya, Spain

**Keywords:** Vestibular assessment, Tail-lift reflex test, Air-righting reflex test, Rat, 3,3’-Iminodipropionitrile

## Abstract

The tail-lift reflex and the air-righting reflex are anti-gravity reflexes in rats that depend on vestibular function. To obtain objective and quantitative measures of performance, we recorded these reflexes with slow motion video in two experiments. In the first experiment, vestibular dysfunction was elicited by acute exposure to 0 (control), 400, 600 or 1000 mg/kg of 3,3’-iminodipropionitrile (IDPN), which causes dose-dependent hair cell degeneration. In the second, rats were exposed to sub-chronic IDPN in the drinking water for 0 (control), 4 or 8 weeks; this causes reversible or irreversible loss of vestibular function depending on exposure time. In the tail-lift test, we obtained the minimum angle defined during the lift and descent maneuver by the nose, the back of the neck and the base of the tail. In the air-righting test, we obtained the time to right the head. We also obtained Vestibular Dysfunction Ratings (VDRs) using a previously validated behavioral test battery. Each measure, VDR, tail-lift angle and air-righting time, demonstrated dose-dependent loss of vestibular function after acute IDPN, and time-dependent loss of vestibular function after sub-chronic IDPN. All measures showed high correlations between each other, and maximal correlation coefficients were found between VDRs and tail-lift angles. In scanning electron microscopy evaluation of the vestibular sensory epithelia, the utricle and the saccule showed diverse pathological outcomes, suggesting that they have a different role in these reflexes. We conclude that these anti-gravity reflexes provide useful objective and quantitative measures of vestibular function in rats that are open to further development.

## INTRODUCTION

The vestibular system includes a diversity of subsystems to cover the existing variety in its natural stimuli, which include linear and angular accelerations of highly diverse magnitudes (Curthoys et al., 2017). It also drives a variety of responses, from vestibulo-ocular and vestibulo-spinal reflexes to endocrine responses and cognitive inputs (Besnard et al., 2015). Therefore, translational studies require the availability of a multiplicity of methods measuring these diverse functions in animals and humans alike (Llorens et al., 2018). At present, a limited number of methods are available for vestibular assessment in rodent, so only part of the vestibular functions can be measured. In addition, many of these methods present technical and practical difficulties that restrain the dissemination of their use or their applicability to certain purposes.

Among the methods historically developed to assess semi-circular canal function in animals, those based on video oculography occupy now a prominent place. These include recording of spontaneous nystagmus (Dyhrfjeld-Johnsen et al., 2013), assessment of post-rotatory nystagmus (Chalansonnet et al., 2018), and direct measure of the vestibulo-ocular reflex (Beraneck et al., 2012; de Jeu and De Zeeuw, 2012; Luebke et al., 2014; Imai et al., 2016). One drawback of these techniques is that they require fixation of the head of the animal. For macular functions, the recording of short-latency vestibular-evoked potentials has been performed in a small number of laboratories (Jones et al., 2011; Sichel et al., 2000). In recent years, the recording of two myogenic potentials evoked by high frequency vestibular stimulation (VEMPs) in the inferior oblique and the sternocleidomastoid muscles, have been demonstrated to assess, respectively, utricular and saccular functions in both humans and laboratory animals (Curthoys et al., 2018). VEMPs are quickly gaining clinical use, but few examples are available of their use in laboratory animals (Vulovic and Curthoys, 2011; Yang et al., 2010; Lo et al., 2015; King et al., 2017).

All the main methods to measure vestibular function cited in the previous paragraph require either to firmly restrain or to anesthetize the animal. Also, they do not cover vestibulo-spinal reflex functions. Loss of vestibulo-spinal input cause gross alterations in spontaneous motor activity and motor reflexes that facilitate the identification of animals suffering vestibular dysfunction due to congenital (Jones and Jones, 2014) or induced (Llorens et al., 1993) causes. However, few studies include quantitative assessment of these alterations. In many cases, the presence or absence of some of these abnormalities is observed or rated to simply corroborate that animals are mutant or bear the lesion (Russell et al., 2003; Wallace et al., 2002). In other cases, a non-specific motor test, such as the rotarod test, is used to evaluate the vestibular deficiency (Schlecker et al., 2011). Precise recordings of spontaneous motor behavior to robustly assess vestibular function have been reported, but they also require the surgical fixation of a device in the skull of the rat (Pasquet et al., 2016). For more than 25 years, our laboratory has been using a behavioral test battery to obtain a semi-quantitative evaluation of vestibular dysfunction in rats and mice exposed to ototoxic chemicals or other vestibular-damaging conditions (Llorens et al., 1993; Llorens and Rodríguez-Farré., 1997; Boadas-Vaello et al., 2005; Soler-Martín et al., 2007; Saldaña-Ruíz et al., 2013; Sedó-Cabezón et al., 2015). This test battery has proven to be reliable, robust and specific to vestibular dysfunction. It has been successfully used in other laboratories (e.g., Luxa et al., 2013), and modified for the assessment of unilateral lesions (Vignaux et al, 2012; Gaboyard-Niay et al., 2016). However, this method is not fully quantitative and relies on subjective assessment. In the present study, we have studied the suitability for objective and quantitative assessment of two anti-gravity reflexes that are part of this behavioral test battery.

The tail-lift reflex and the air-righting reflex are widely recognized to depend on vestibular function. The tail-lift test (Hunt et al., 1987; Llorens et al., 1993) evaluates the loss of the anti-gravity body and limb extension response. The air-right test (Ossenkopp et al., 1990) evaluates the loss of the anti-gravity righting-in-the-air response. In the present study, we evaluated whether these two reflexes could be used for quantitative assessment by high-speed video recording and subsequent movie analysis. We compared the result of these analyses with that of our previously established semi-quantitative behavioral test battery, using control rats and rats suffering vestibular dysfunction following ototoxic exposure. The data obtained demonstrate the suitability of this non-invasive approach to measure vestibular dysfunction in awake rats.

## MATERIAL AND METHODS

### Animals and treatments

This study used two lots of male Long-Evans rats, aged 8-9 weeks, purchased from Janvier Labs (Le-Genest-Saint-Isle, France). The rats were housed in groups of three in standard cages (215 × 465 × 145 mm) with a bedding of wood shavings. They were acclimatized for six days before experimentation, were maintained on a 12:12 h light:dark cycle (07: 30-19:30 h) at 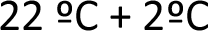, and had free access to standard diet pellets (TEKLAD 2014, Harlan Laboratories, Sant Feliu de Codines, Spain). On a regular basis, the animals were weighed and evaluated for overall toxicity to limit suffering according to ethical criteria. The use of the animals was in accordance with EU Directive 2010/63 as implemented by Law 5/1995 and Act 214/1997 of the Generalitat de Catalunya, and Law 6/2013 and Act 53/2013 of the Gobierno de España. The experiments were approved by the Ethics Committee on Animal Experimentation of the Universitat de Barcelona.

The study included two experiments evaluating the effects of acute and sub-chronic exposure to IDPN. In the acute exposure experiment, rats were administered 0 (Control group), 400 (IDPN 400), 600 (IDPN 600), or 1000 (IDPN 1000) mg/kg of IDPN (>98%, TCI Europe, Zwijndrecht, Belgium), i.p., in 2 ml/kg of saline (n=6 / group), in accordance with published data (Llorens et al., 1993). The vestibular function of the animals was evaluated before administration, and at nine time-points up to 13 weeks after exposure. In the sub-chronic exposure experiment, IDPN was administered in the drinking water. Rats were exposed to tap water only (Control group), or to water containing 20 mM IDPN for 4 (IDPN 4-wk) or 8 (IDPN 8-wk) weeks (n=9 / group). These animals were evaluated for vestibular function before exposure and at weekly or two-week intervals for 16 weeks.

At the end of the experimental periods, rats were given an overdose of chloral hydrate (800 mg/kg, i.p.) and decapitated under deep anesthesia for collection of the vestibular sensory epithelia. The temporal bones were immersed in cold fixative for immediate collection of the vestibular sensory epithelia under a fume hood. The first ear was dissected in 2.5% glutaraldehyde in 0.1 M cacodylate buffer (pH 7.2) and the epithelia were processed for scanning electron microscopy (SEM) observation of the hair bundles in their surface. The second ear was collected for studies not included in the present work.

### Assessment of vestibular dysfunction

The loss of vestibular function was assessed by three means. First, a measure was obtained using a semi-quantitative behavioral test battery that has been proven to be sensitive and specific to vestibular function, as reported previously (Llorens et al., 1993; Llorens and Rodríguez-Farré, 1997; Boadas-Vaello et al., 2005; Sedó-Cabezón et al., 2015). In this battery, 6 items are rated from 0 (normal behavior) to 4 (highest score of behavioral deficiency) to obtain a sum score of 0 to 24 (Vestibular Dysfunction Rating, VDR). Three of these items are alterations in spontaneous behavior (stereotyped circling, retropulsion, and head bobbing) that appear in vestibular-deficient animals. The other three are alterations in reflexes: tail-lift reflex, contact inhibition of the righting reflex, and air-righting reflex. More complete descriptions of this battery have been published previously (Llorens et al., 1993; Boadas-Vaello et al., 2005). In addition, two of the tests included in this battery were recorded by high speed video for slow-motion analysis of the rat’s reflexes. In the air-right reflex tests, the rat is held in supine position at approximately 40 cm of height and released to fall on a foam cushion. In the tail-lift reflex test, the rat is grasped by the base of the tail, gently lifted to approximately 40 cm and returned down.

Motor reflexes were recorded using a Casio Exilim ZR700 camera, at 240 frames per second (fps), and 512 x 384 pixels. The tail-lift reflex was recorded from the rat’s side, while the air-righting reflex was recorded from the front side. To facilitate video tracking, the tail-lift reflex was recorded on red background and, in the chronic experiment, a white marble was placed with a small rubber band collar in each animal as back neck marker. Video analysis was performed using the free software Kinovea (www.kinovea.org). For the air-righting reflex, the stopwatch element was used to obtain the time from the release of the animal until it fully righted its head, in the air or already on the foam. For the tail-lift reflex, each rat was tracked from the nose, neck and base of the tail through the whole movement in a semi-automatic manner. The locations of the points are computed automatically, but can be adjusted at any time if necessary. For further analysis, the data generated by the tracking process were exported to text files.

R programming language was used to analyze the data obtained from the tail-lift reflex. Mathematical calculations of the script for angular changes were performed using the coordinates (Fig. 1) as a function of time obtained from Kinovea. The dot product of the two vectors a and b is defined by

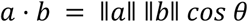
where *θ* is the angle between *a* and *b* (radiants were converted into degrees). Therefore, the angle between nose, neck and tail was calculated as:

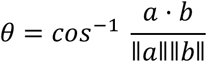
where *a* · *b* is the dot product of the two given vectors. The nose-neck-tail angle was determined as a function of time, and the minimum angle shown by the rat during the tail-lift movement was calculated. Further details of the procedure and the R script are provided in the Annexes.

**Figure 1.**
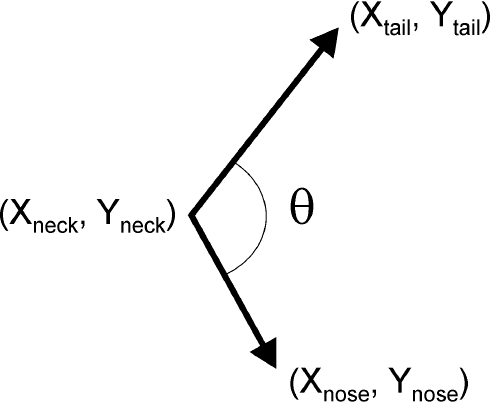
Geometric interpretation of the angle between vector the tail-to-neck vector and the neck-to-nose vector. Analysis of the video movies provide the X and Y coordinates of nose, neck and tail marks at 1/240 s intervals.

### Scanning Electron Microscopy

After dissection, the vestibular epithelia were fixed overnight at 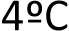. They were then post-fixed for 1 h in 1% osmium tetroxide in cacodylate buffer, transferred to 70 % alcohol, and stored at 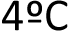 until further processing. Epithelia were then dehydrated with increasing concentrations of ethanol, up to 100%, dried in a critical-point dryer using liquid CO2, coated with carbon, and observed in a JEOL JSM-7001F field emission scanning electron microscope at 15 kV. To assess hair cell loss, the number of hair bundles was obtained in images at 1,500X magnification from the striolar region.

### Statistics

Data are shown as mean +/− SE. They were analyzed by repeated measures MANOVA (Wilk’s criterion) with day as the within-subject factor. Day-by-day comparisons were performed using one-way ANOVA followed by Duncan’s post-hoc tests. The Pearson coefficient was used to evaluate the correlation of pairs of data from the same animals. All analyses were done with the IBM SPSS 20 program package.

## RESULTS

### Effects of IDPN on body weight and overall health

Acute and chronic exposure of male adult Long-Evans rats to IDPN caused effects on body weight matching the data reported previously (Llorens et al., 1993; Llorens and Rodríguez-Farré, 1997; Sedó-Cabezón et al., 2015). Acute exposure to 400 mg/kg IDPN caused no significant effects on body weight, while exposure to 600 or 1000 mg/kg caused a decrease in body weight that progressed for 4-5 days, after which animals resumed body weight gain at a similar rate as control animals. The rats in these two high-dose groups displayed the overt alterations in spontaneous behavior that characterize vestibular deficient animals (Llorens et al., 1993), but showed no other signs of systemic toxicity, and all of them survived the entire experimental period of 91 days. Animals given chronic IDPN by drinking water exposure ceased increasing their body weight as control animals do. Their body weights restarted monotonous increase after the end of the exposure. At weeks 11-12, one rat exposed to IDPN for 8 weeks was euthanized according to ethics protocol due to unexpected appearance of paraparesis, and one rat exposed for 4 weeks was found death, likely due to fighting with its cage-mates.

### Effects of acute IDPN on measures of vestibular function

Rats exposed to acute IDPN showed a dose-dependent loss of vestibular function as shown by VDR data (Fig. 2A). Thus, IDPN 400 animals showed a very small and transient effect on VDR, while IDPN 600 and IDPN 1000 animals showed high VDRs shortly after exposure and stably until the end of the experiment. Maximal effect was evident in IDPN 1000 rats already at day 3 after exposure, but only by day 7 in IDPN 600 rats. MANOVA analysis resulted in significant effects of the day factor (F(9,12)=156.2, p<0.001), treatment factor (F(3,20)=729.8, p<0.001), and day-by-treatment interaction (F(27, 35.7)=16.6, p<0.001). Day-by-day ANOVA analysis resulted in significant differences between treatment groups, at all days from day 3 to day 91 (all F(3,20)>97.7, all p<0.001).

**Figure 2.**
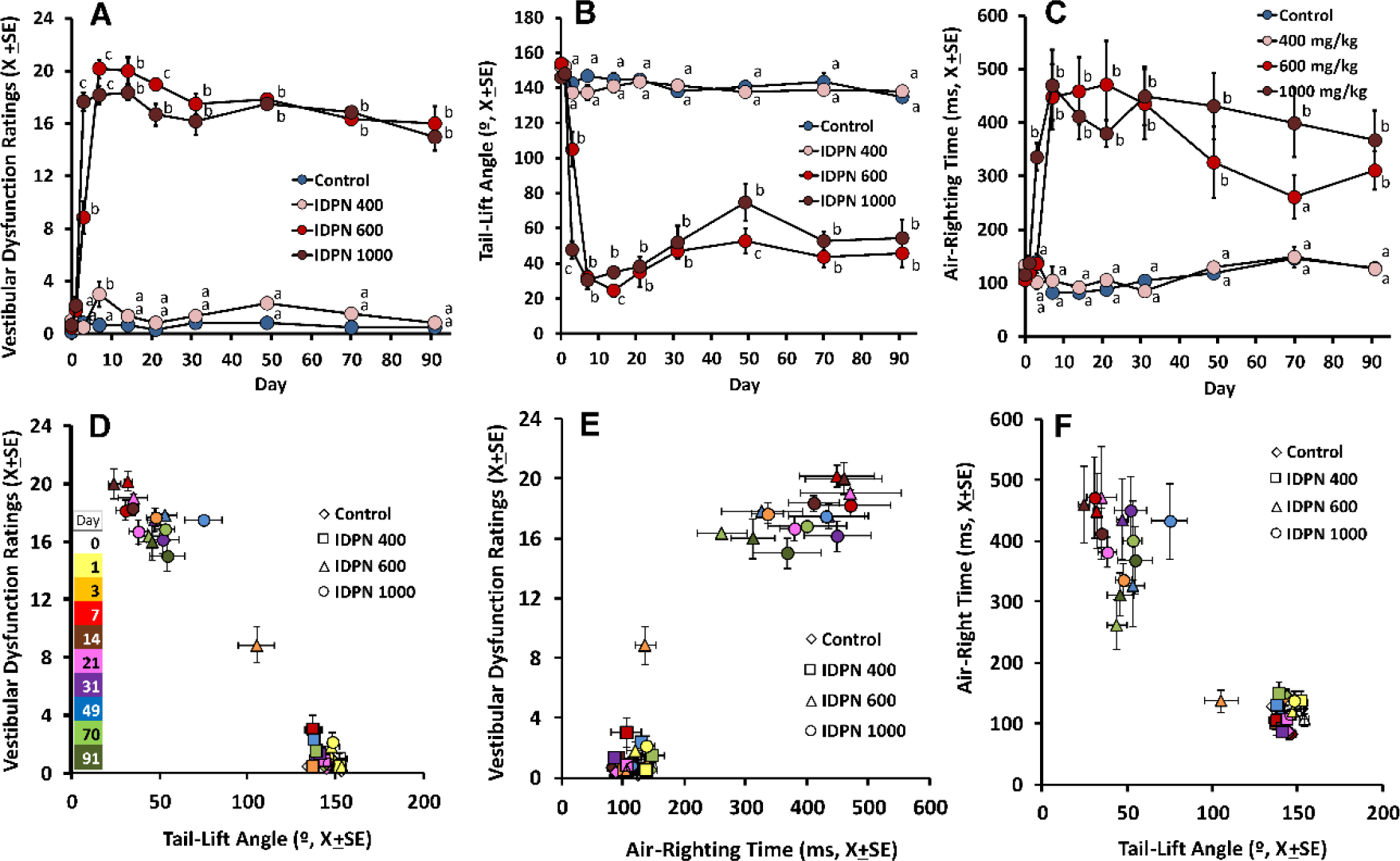
Effects of acute IDPN on vestibular function. Data are X+SE (n=6/group) of rats treated with IDPN at 0 (control), 400, 600 or 1000 mg/kg. a, b, c: groups signaled with different letters are significantly different from each other, P<0.05, Duncan’s test after significant ANOVA at that day. A) Vestibular Dysfunction Ratings (VDRs). Data are sum ratings from a battery of 6 tests sensitive to vestibular dysfunction, each rated 0 (normal behavior) to 4 (extreme evidence of vestibular loss). B) Tail-lift angle. Data are minimum nose-neck-base of the tail angles shown by the rats when lifted by the tail and returned down. C) Air-righting time. Data are times elapsed from the moment when the rat is released supine in the air and the moment it rights its head. D, E, F) Pair comparisons with VDRs, tail-lift angles and air-righting times. Each point corresponds to a treatment group (identified by shapes) and time points (identified by colors as shown in panel D).

The loss of vestibular function in IDPN 600 and IDPN 1000 rats was also shown by a decrease in the minimum nose-neck-tail angle attained by the animal during the tail-lift test (Fig. 2B). Control rats typically showed angles between 130 and 160 °, and these tail-lift angles were not modified in the IDPN 400 group. However, minimum angles were largely reduced in IDPN 600 and IDPN 1000 rats, which curled instead of showing the normal extension response for landing. The reduction in tail-lift angles was persistent and similar in both these high-dose groups. The only relevant difference between these two groups was found in the velocity at which the loss of function appeared: in the IDPN 1000 group it was maximal already at day 3 after injection, while in the IDPN 600 group this effect was not full yet at that time point. MANOVA analysis resulted in significant effects of the day factor (F(9,12)=343.9, p<0.001), treatment factor (F(3,20)=348.3, p<0.001), and day-by-treatment interaction (F(27, 35.7)=15.8, p<0.001). Day-by-day ANOVA analysis resulted in significant differences between treatment groups, at all days from day 3 to day 91 (all F(3,20)>43.8, all p<0.001).

In the air-righting test (Fig. 2C), the vestibular toxicity of IDPN resulted in an increase in the time to right in the IDPN 600 and IDPN 1000 groups, but not in the IDPN 400 group. At day 3, a significant increase in time was recorded in the IDPN 1000 but not in the IDPN 600 animals. At day 7, both high-dose groups attained maximal values. At long times after dosing, the air-righting times in these groups tended to decrease, but did not attain control-like values. MANOVA analysis resulted in significant effects of the day factor (F(9,12)=61.8, p<0.001), treatment factor (F(3,20)=95.8, p<0.001), and day-by-treatment interaction (F(27, 35.7)=15.6, p<0.001). Day-by-day ANOVA analysis resulted in significant differences between treatment groups, at all days from day 3 to day 91 (all F(3,20)>9.2, all p<0.001).

Figure 2 also shows the relationship between the three measures of vestibular function, depicted as mean and error values for all groups and time points. When all individual and time values (n=240) were used to calculate the Pearson correlation coefficient, a very high coefficient was obtained between VDR and tail-lift angle (−0.957, p<0.001). High but lower correlation coefficients were obtained between VDR and air-righting time (0.848, p<0.001), and between tail-lift angle and air-righting time (−0.819, p<0.001).

### Effects of sub-chronic IDPN on measures of vestibular function

Rats exposed to 20 mM of IDPN suffered a progressive loss of vestibular function as revealed by VDR data (Fig. 3A). The effect was first noticed at 2 weeks of exposure and increased largely during the next two weeks. The animals exposed for 4 weeks showed a significant recovery after termination of the exposure, and group mean values were not different from control group means at the end of the survival period. The functional recovery occurred mainly in the three initial weeks after the end of the exposure. In the group of rats exposed for 8 weeks, the loss of function showed some additional progression between 4 and 8 weeks. In contrast to what observed in the IDPN 4-wk group, termination of the exposure did not trigger significant functional recovery in the IDPN 8-wk rats, and the animals showed high VDRs until the end of the experiment after a recovery period of 8 weeks. MANOVA analysis included weeks 0 to 11, because two animals were lost at week 12 (see above). The analysis resulted in significant effects of the day factor (F(11,14)=30.9, p<0.001), treatment factor (F(2,24)=42.7, p<0.001), and day-by-treatment interaction (F(22, 28)=10.6, p<0.001). Day-by-day ANOVA analysis resulted in significant differences between treatment groups, at all weeks from week 3 to week 16 (all F(2,24) or F(2,22) >11.2, all p≤0.001).

**Figure 3.**
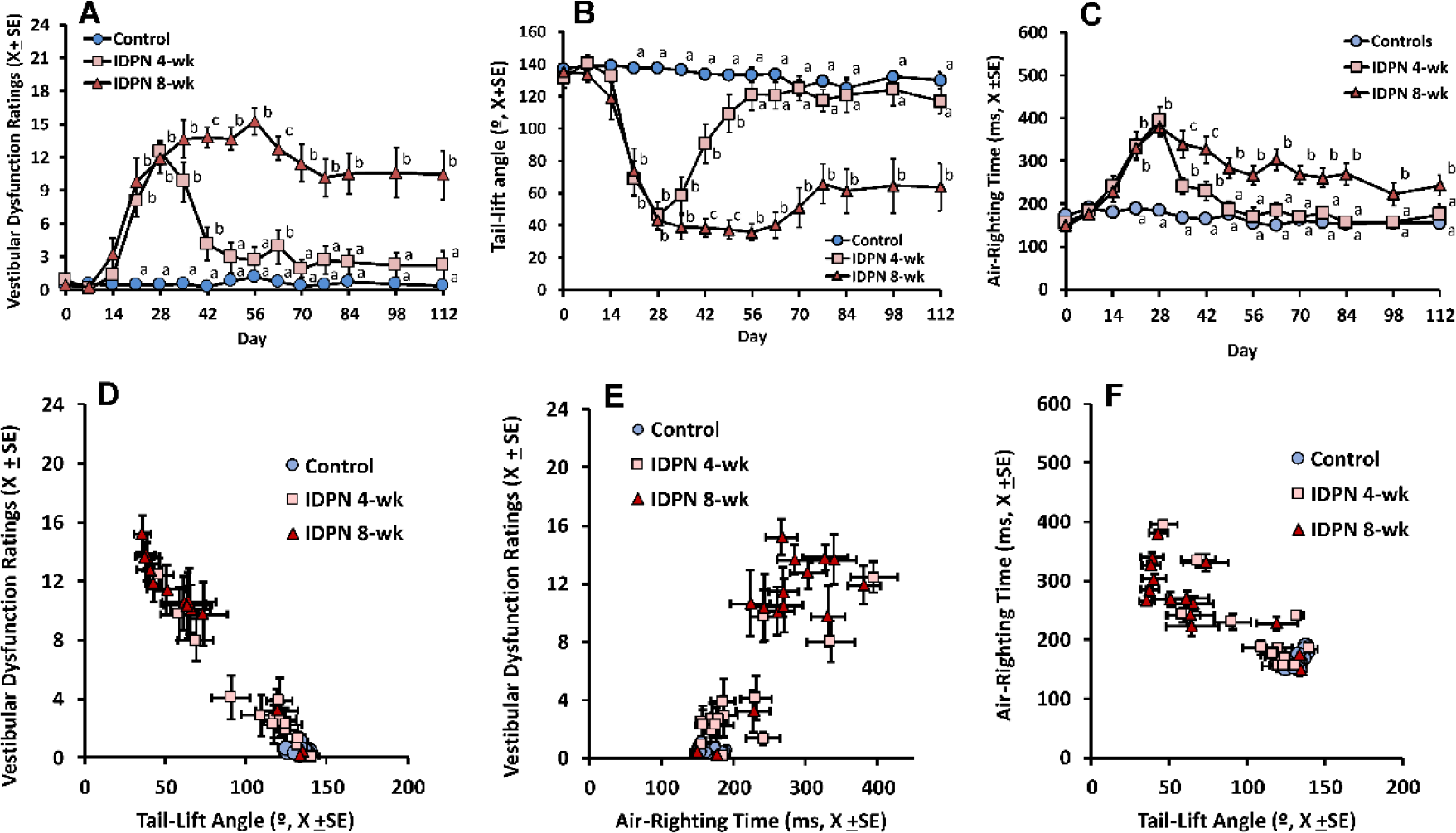
Effects of sub-chronic IDPN on vestibular function. Data are X+SE (n= 8-9/group) of rats treated with 20 mM of IDPN in the drinking water for 0 (control), 4 or 8 weeks. a, b, c: groups signaled with different letters are significantly different from each other, P<0.05, Duncan’s test after significant ANOVA at that day. A) Vestibular Dysfunction Ratings (VDRs). Data are sum ratings from a battery of 6 tests sensitive to vestibular dysfunction, each rated 0 (normal behavior) to 4 (extreme evidence of vestibular loss). B) Tail-lift angle. Data are minimum nose-neck-base of the tail angles shown by the rats when lifted by the tail and returned down. C) Air-righting time. Data are times elapsed from the moment when the rat is released supine in the air and the moment it rights its head. D, E, F) Pair comparisons with VDRs, tail-lift angles and air-righting times. Each point corresponds to a treatment group (identified by shapes and colors) and time points.

The loss of vestibular function in rats exposed to sub-chronic IDPN was also shown by a decrease in the minimum nose-neck-tail angle attained by the animal during the tail-lift test (Fig. 3B). Minimum angles were largely reduced by 3 weeks of exposure, and further decreased at 4 weeks. After the 4-week time point, a complete recovery occurred in the group of rats in which the exposure was terminated. In contrast, the mean minimum angles remained low in the animals exposed for 8 weeks for the whole exposure period, and showed little recovery after the end of the exposure. MANOVA analysis for weeks 0 to 11 resulted in significant effects of the day factor (F(11,14)=20.1, p<0.001), treatment factor (F(2,24)=48.3, p<0.001), and day-by-treatment interaction (F(22, 28)=8.1, p<0.001). Day-by-day ANOVA analysis resulted in significant differences between treatment groups, at all weeks from week 3 to week 16 (all F(2,24) or F(2,22) >10.8, all p<0.001).

In the air-righting test (Fig. 3C), the IDPN 4-wk rats showed an increase in righting times that progressed during exposure, and was followed by complete recovery in mean group values after the end of the exposure. In the rats exposed to 20 mM IDPN for 8 weeks, maximal effects on air-righting times were observed at 4 weeks of exposure. After this time point, a slowly decline in times was observed, but this decline was not accelerated after the end of the exposure at 8 weeks, and air-righting times were significantly increased in this group in comparison to control and IDPN 4-wk animals until the end of the experiment. MANOVA analysis for weeks 0 to 11 resulted in significant effects of the day factor (F(11,14)=18.9, p<0.001), treatment factor (F(2,24)=30.1, p<0.001), and day-by-treatment interaction (F(22, 28)=6.9, p<0.001). Day-by-day ANOVA analysis resulted in significant differences between treatment groups, at all weeks from week 3 to week 16 (all F(2,24) or F(2,22) >5.6, all p≤0.01).

Figure 3 also shows the relationship between the three measures of vestibular function in the sub-chronic IDPN experiment, depicted as mean and error values for all groups and time points. Correlation coefficients calculated on all individual and time values (n=399) were: −0.937 between VDR and tail-lift angle (p<0.001), 0.764 between VDR and air-righting time (p<0.001), and −0.741 between tail-lift angle and air-righting time (p<0.001).

Examples of individual values of the tail-lift angles of rats receiving sub-chronic IDPN are shown in Figure 4. These data illustrate the complete recovery of some animals exposed for 4 weeks even after attaining a deep loss of function at the end of the exposure period, and the incomplete recovery of the worst case example (Fig. 4A). In animals exposed for 8 weeks (Fig. 4B) good functional recovery was still recorded in some cases, but a complete failure of recovery was recorded in the worst cases.

**Figure 4.**
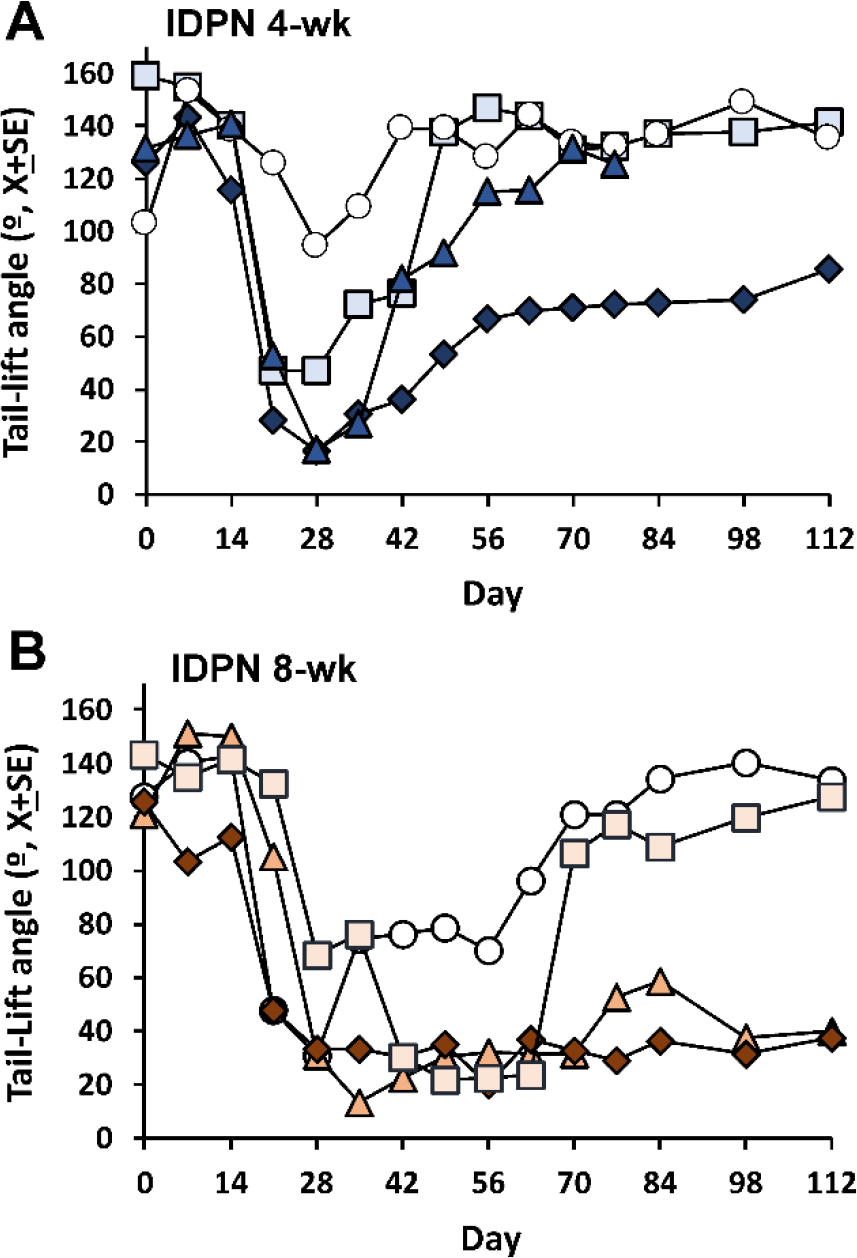
Effects of sub-chronic IDPN on tail-lift angles. Data are from representative individual animals exposed to 20 mM of IDPN in the drinking water for 4 (A) or 8 (B) weeks.

### Effects of IDPN on surface morphology of the vestibular sensory epithelia

SEM analysis of the vestibular epithelia surfaces revealed a dose-dependent loss of hair bundles after acute IDPN (Fig. 5). The progression of the damage showed a crista > utricle > saccule gradient. In the crista (not shown) and the utricle (Fig. 5A-D), loss of hair bundles was evident in the central or peri-striolar regions of some IDPN 400 rats; the loss was almost complete or complete in all IDPN 600 and IDPN 1000 rats. In the saccule (Fig. 5E-H), the extent of the damage in the IDPN 600 rats was lesser than in the utricle. Hair bundle counts in the utricle (Fig. 5I) demonstrated significant group differences (F(3,17)=48.5, p<0.001)). No significant difference was recorded between Control and IDPN 400 rats, and a significant loss of hair bundles was demonstrated in rats exposed to 600 or 1000 mg/kg of IDPN. In the saccule (Fig. 5L), significant group differences (F(3,15)=41.6, p<0.001)) resulted from a 40% loss of stereocilia bundles in IDPN 600 animals and an almost complete (99%) loss in IDPN 1000 animals.

**Figure 5.**
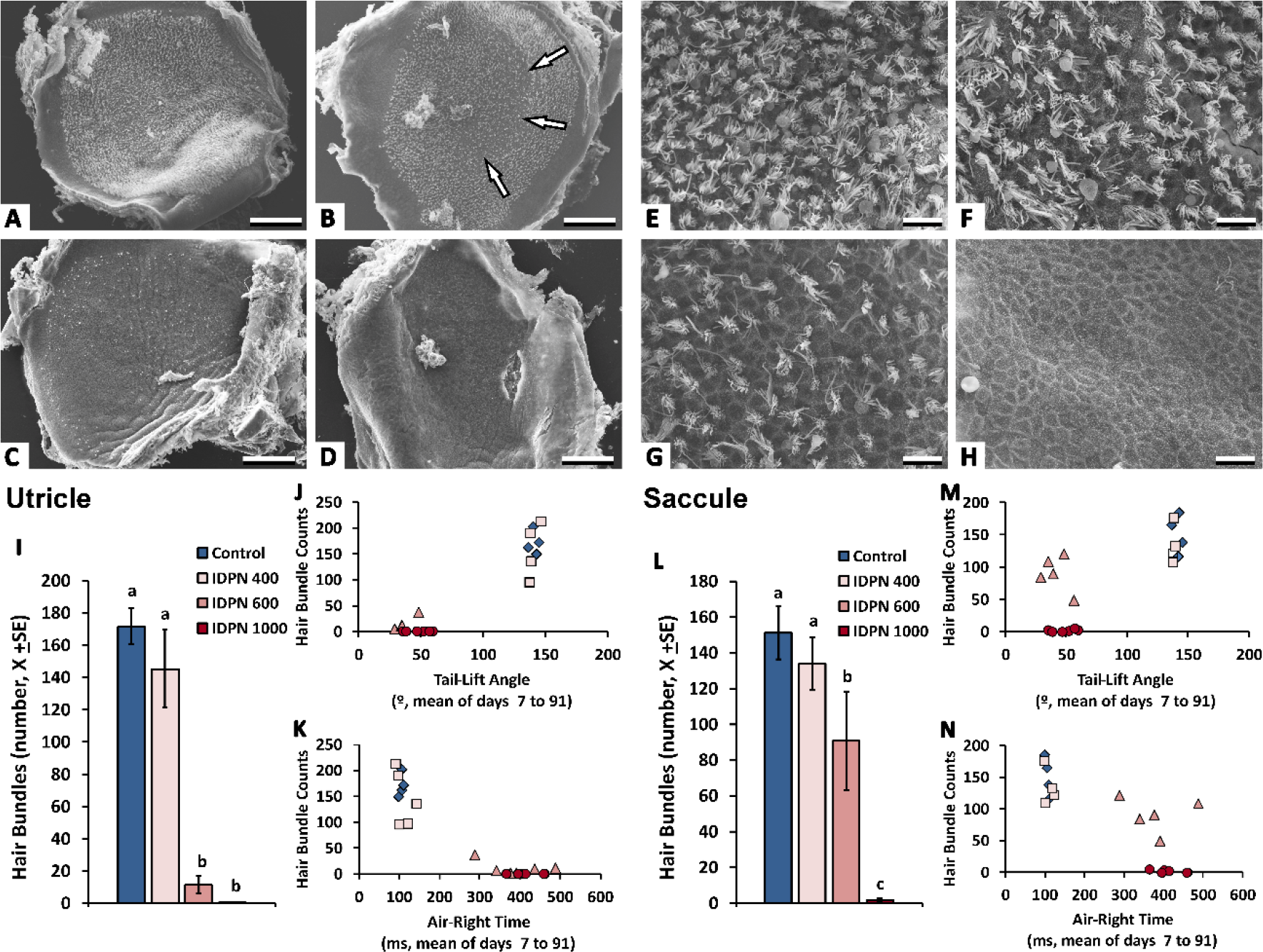
Effect of acute IDPN on the utricle and the saccule as assessed by SEM analysis of the surfaces of the sensory epithelia. Upper panels show surface images from utricules (A-D) and the central (peri-striolar) region of saccules (E-F) from animals exposed to 0 (control, A and E), 400 (B and F), 600 (C and G), and 1000 (D and H) mg/kg of IDPN. Arrows in B indicate the peri-striolar zone. Scale bars: 100 μm in A-D; 10 μm in E-H. (I and L) Number of hair bundles in 1,500X images of the peri-striolar region of utricles and saccules, respectively. a, b, c: groups signaled with different letters are significantly different from each other, P<0.05, Duncan’s test after significant ANOVA. (J, K, M and N) Relationship between hair bundle counts in the utricle (J and K) and saccule (M and N), and tail-lift angles (J and M) and air-righting times (K and N). Each point corresponds to the paired data of a single animal, exposed to 0 (control, blue diamonds), 400 (squares), 600 (triangles) or 1000 (circles) mg/kg of IDPN.

Figure 5 also shows the relationship between hair bundle counts in the utricle (Fig. 5J, 5K) and the saccule (Fig. 5M, 5N) with the tail-lift angle (Fig. 5J, 5M) and the air-righting time (Fig. 5K, 5N). Significant correlations were found between these quantitative measures of vestibular pathology and vestibular function, although these arise mostly from the all or none effects differentiating the Control and IDPN 400 groups from the IDPN 600 and IDPN 1000 groups. One exception was the intermediate bundle counts in the saccule of the IDPN 600 animals. The highest correlation coefficient was found for the number of hair bundles in the utricle and the tail-lift angle (0.863, p<0.001, n=21), followed by that of the utricle counts and the air-righting time (−0.754, p<0.001, n=21). Saccular counts had lower correlation coefficients with the tail-lift angle (0.685, p=0.001, n=19) and the air-righting time (−0.659, p=0.002, n=19), as IDPN 600 animals showed maximal behavioral effects with only partial loss of hair bundles.

Most of the SEM samples from the sub-chronic IDPN experiment were lost due to a failure in the critical-drying process, so this experiment was not available for quantitative analysis. Assessment of the remaining samples indicated the existence of a link between persistent loss of function and loss of hair bundles (not shown).

## DISCUSSION

The design of new methods to evaluate vestibular function faces several difficulties. One is that the adequate stimuli for vestibular stimulation are angular and lineal accelerations of the head, which are not easy to deliver to experimental animals (Jones et al., 2011; Beraneck et al., 2012; de Jeu and De Zeeuw, 2012). Another difficulty is that the endpoints to be measured are usually indirect measures of vestibular function, motor responses that are controlled by vestibular function, but that receive also inputs from other systems (Serra et al., 2013; Allum and Carpenter, 2013). Although direct recording of vestibular signals can be envisaged, electrical potentials from the vestibular periphery are small, as are the vestibular-evoked potentials (Brown et al., 2017). In addition, cortical areas receiving vestibular input are poorly defined and widely distributed (Brown et al., 2017). In the present study, we have studied the suitability for quantitative assessment of two anti-gravity reflexes for vestibular assessment in rats. These reflexes rely on the constant gravitational acceleration as stimulus, and are therefore based on a completely stable trigger cause. Also, they consist in highly stereotyped motor responses, on which the visual system may have little impact.

Performance of the tests and obtention of movies suitable for the analysis require some practice, but do not pose major difficulties. For instance, rats may turn the body to one side when lifted by the tail, this resulting in a movie offering a ventral or dorsal view of the animal instead of the adequate side view. When this happened, the test was repeated a second or third time, and the best movie used for the analysis. Using video recording at 240 fps we obtained movies suitable to calculate the minimum angle in the tail-lift test and the time to right in the air-righting test. This speed of recording is only 8 times the standard speed of video recording (30 fps) and is nowadays available in many domestic photography and video cameras, including smartphones. For the tail-lift test, we explored the utility of alternate measures, such as the mean angle during a certain period around the peak of the reflex and the nose-tail velocity difference, but these resulted in higher within-group variabilities and did not provide any apparent advantage respect to the minimum angle (data not shown). However, the possibility that these tests may provide additional meaningful measures of vestibular function remains open.

To validate the utility of the tests, we used the acute and sub-chronic IDPN models. Acute IDPN cause a dose-dependent degeneration of hair cells resulting in the fast loss of vestibular function, which persists throughout the experimental period (Llorens et al., 1993; Llorens and Demêmes, 1994; Seoane et al., 2001b; Soler-Martín et al., 2007). In contrast, sub-chronic exposure results in a slowly evolving loss of vestibular function (Llorens and Rodríguez-Farré, 1997) that associates with hair cell extrusion (Seoane et al., 2001a; Seoane et al., 2001b), preceded by a first phase characterized by reversible synaptic uncoupling and hair cell detachment (Sedó-Cabezón et al., 2015; Greguske et al., 2019). Both the tail-lift angle and the air-right time revealed the persistent nature of the deficits caused by acute IDPN and the potential reversibility of the deficits caused by sub-chronic IDPN. Comparison of the high concordance between the VDR graphs and the tail-lift angle graphs, and consideration of the high correlations recorded between these two measures indicate that the tail-lift angle is a new objective and quantitative measure likely measuring the same functional alteration than the VDR test battery. It is worth noting that the none-to-high effect jump observed between the IDPN 400 and the IDPN 600 animals in VRD scores is corroborated by the tail-lift angles. The dose of 400 mg/kg of IDPN was selected to have an intermediate effect according to previous data (Llorens et al., 1993), but it failed to cause a significant impact on vestibular function in this study. In fact, although hair cell loss was observed in some animals of this group, others presented intact vestibular epithelia and no statistically significant loss of hair cells was recorded in this group in either the utricle or the saccule.

Compared to the VDRs and the tail-lift angles, the air-righting time showed higher variabilities, an apparent higher tendency to recovery in the most affected groups, and smaller correlation coefficients with the other two measures. An explanation for these differences would be that the air-righting reflex depends on functional inputs partially different from those on which the tail-lift reflex depends, or may be more prone to compensation through learning or use of information from other sensory systems. Future investigations may provide answer to these hypotheses.

As a pilot investigation on the cellular basis of these reflexes, we planned the quantification of hair cell loss using SEM assessment of hair bundle density in the maculae. Most samples from the sub-chronic experiment were lost, but those of the acute experiment allowed for the comparison of the results of the reflex tests with hair cell counts in the utricule and the saccule. For both reflexes, animals fell in one of two groups that were coincidental with dose groups and hair cell counts in the utricle. On one side, control and IDPN 400 animals showed normal or almost normal behaviours and high densities of stereocilia bundles in the utricle. On the other side, IDPN 600 and IDPN 1000 animals showed deeply altered behaviours and complete or almost complete loss of utricular bundles. At difference, saccular counts for IDPN 600 animals did not fit this pattern, because those animals showed deep behavioural dysfunction despite preservation of a significant proportion of stereocilia bundles in the saccule. These data suggest that these reflexes may relate more to utricular than saccular function. While these data are clearly insufficient to make conclusions on the cellular basis of the tail-lift and the air-right reflexes, they indicate that the tail-lift angle and the air-righting time may eventually become measures of specific vestibular functions with well-characterized anatomical substrates.

In conclusion, the present study demonstrates the suitability of high-speed video recording of two anti-gravity reflexes to obtain objective and quantitative measures of vestibular dysfunction in rats. The data obtained demonstrate that the minimum angle drawn by the nose, the neck and the base of the tail when the rat is lifted by the tail, and the time to right the head in the air-right reflex test provide good measures of permanent and transient loss of vestibular function, as elicited by acute or chronic exposure to IDPN. These measures appear as reliable and easy to implement and appear suitable for further development.

## Acknowledgements

This study was supported by grants BFU2015-66109-R (Ministerio de Economia y Competitividad, MINECO/FEDER, EU), and 2017 SGR 621 (Agència de Gestió d’Ajuts Universitaris i de Recerca, Generalitat de Catalunya). E.A.G. was supported by the Secretaria d’Universitats i Recerca del Departament d’Economia i Coneixement de la Generalitat de Catalunya (FI-DGR 2015 Program) and by the Ministerio de Educación, Cultura y Deporte de España (FPU 2015). The scanning electron microscopy studies were performed at the Scientific and Technological Centers of the University of Barcelona (CCiT-UB). We thank Josep M. Rebled and Eva Parts for technical assistance. We also thank Meritxell Deulofeu, Sílvia Prades and Adrià Ricarte for their contributions to the study as part of their final degree projects.

## APPENDIX A Evaluation of the tail-lift and air-righting video recordings using Kinovea and R language

This section describes the analysis of the tail-lift and air-righting reflexes using the Kinovea software (www.kinovea.com).

## 1. Kinovea main settings

In the main menu, enter the number of seconds per frame at capture time, as the capture speed (240 fps) differs from the default standard speed (30 fps).

Use the calibrate tool to establish the relationship between pixels and real distances. Draw a line on the image, and enter the real length of the segment. The line can be deleted afterwards to avoid disturbing the video analysis, yet the calibrate remains as determined.

Select a point as origin of the coordinate system that will be used.

Select preferred time representation.

## 2. Analysis of the tail-lift reflex

Each rat is tracked from the nose, neck and tail through the whole movement. To start, select a cross marker from the drawing tool-bar and place them on the nose, neck (white marble) and on the base of the tail.Use this order, as this is the order for which the analysis script is written.

Right click on the cross marker and select Track path. Move the video forward by using the Play button. For more complex tracking, use the Next Frame button or the Mouse Wheel to forward the video step by step. In Kinovea, tracking is a semi-automatic process. The locations of the points are computed automatically, yet if needed they can be adjusted at any time.During the tracking process, disoriented points can be adjusted back to their correct positions by the outer rectangle. Drag from the outer rectangle until the cross at the centre of the tracking tool is at its correct location. To finish tracking, right click and select End Path Edition. In case some points are still misplaced, the path can be edited again by right clicking the path and selecting Restart Path Edition. The path can be drawn in various ways and this might be useful when saving the video. Display options can be accessed by right clicking the Path track and selecting Configuration. Additionally, distance and speed measurements are available in the configuration.

After finishing the tracking, the video can be saved by including solely the frames of interest. To this, click on Start Working Zone and End Working Zone.

There are multiple options to save the video. Choosing “*Combine video and key images data in the file*” generates a file that can be opened again in Kinovea, to modify the tracked path and export the data again, if needed. Choosing “*permanently paint key images data on the video*” will generate a final video of the tracked path that cannot be further modified but can be open with any video player.

For subsequent analysis, in the main menu select Export to Spreadsheet > Trajectories to simple text.

Before analyzing a second rat, the paths tracked for the first rat must be deleted if it is on the same video, because the new tracking do not override the previous tracking. This problem can be avoided if the videos are made separately for each rat.

## 3. Analysis of the air-righting reflex

Select a stopwatch from the drawing tool-bar and place it on the image by clicking a desired location.

Activate the stopwatch when the rat is released by the experimenter, and stop the stopwatch when the head of the rat is fully turned into the upright position. Save the video choosing the option of “*permanently paint key images data on the video*”.

## 4. Processing of tail-lift data

To enable processing with the R script included as Annex B, the text file must be named as follows: Timepoint_X_”Cage-Number”_”Rat-ID”, where X is the time-point number. For example, Week_3_R00950_LF can be used for naming. If the name misses the underscores, the R script cannot read the file.

Next, the obtained data files must be ordered into subfolders. Create a main folder where you are going to place all the data files. The folder can be named as desired, yet it must follow the rules of R. It cannot contain any spaces neither special marks. We used “Coordinates” for our folder name. Then create folders for each week separately after processing the data of the time point. Name them as follow: Timepoint_X (for example Week_10 or Day_10). For each time point folder, place new folders for each treatment group separately. For instance, you can have three experimental groups named Control, Treatment_1 and Treatment_2. Now when you have tracked the rats, place them into corresponding folder. The name of the folder will be seen in the titles of the graphs.

After saving the data file, open it and make sure that all the setups were correctly adjusted and the file contains data only form one rat. When the data file is opened, all the tracked coordinates for each marker can be seen one below another, in order: nose, neck and tail. The left column shows the time measured for the tracking. Since we chose total milliseconds for the time representation, the whole movement should be around 400 to 800 ms. If it shows oddly great numbers, like 1000 – 8000 ms, make sure that you have adjusted the time display to 240 fps. Next columns contain the X and Y positions, respectively. From the Y position, the order of the markers can be verified. For example, if it is a control rat, the Y position of the nose should start at the lowest point, then should come the neck and lastly the tail. More importantly, verify that each of the markers were tracked the same amount of time. If the columns are unequal length, there will be an error when processing the data files.

Finally, after opening the script (Appendix B) with R (we used RStudio as a development environment for R), the main folder path must be set as working directory in the script. Meaning that the script knows to search the data files from the main file set earlier. Copy and paste the directory on full_path = “Set here the address of your main folder”. Remember to add slash at the end of the path.

~~~
*# Always add at the end of the path*
full**_path** =
”/Users/vanessamartinslopes/Documents/Master_thesisE0510_Vanessa_IDPN_Chron ic_in_ratCoordinates/”
~~~

~~~
*# Read all TXT files on directory*
**list_**weeks **<**- **list**.dirs (full**_path**, full. **names** = FALSE, recursive = FALSE)
**for** ( | in 1 : **length** ( **list_**weeks)) {
run_script ( **paste** ( full**_path**, **list _**weeks [|], “/”, sep= “”))
}
~~~

## APPENDIX B R Script used to analyse the nose-neck-tail track data generated by the Kimovea software from the tail-lift video recordings.

~~~
**cat** (“Starting␣Script”)
**cat** (“Cleaning␣variables”)
**rm** (**list** = **ls**())
~~~

~~~
**library** (ggplot2)
**library** (reshape2)
**library** (scales)
**library** (gridExtra)
**library** (cowplot)
**library** (xlsx)
~~~

~~~
**cat** (“Libraries␣loaded”)
~~~

~~~
*# Function to process data*
process**_data_**f **<**− **function**(**df**) {
**df** <− **do.call**(**data.frame**, **aggregate**(. ~V1, **df**, **as.vector**))
**df** <− setNames(**df**, **c**(“time”,“xnose”,“xneck”,“xtail”,“ynose”,“yneck”,“ytail”))
~~~

~~~
*# (a1, a2), (b1, b2) and (y1, y2) points of the vectors*
dfpoints **<**− **data.frame**(**df$**xneck−**df$**xtail, **df$**yneck−**df$**ytail, **df$**xneck−**df$**xnose,
**df$**yneck−**df$**ynose, **df$**xtail−**df$**xnose, **df$**ytail−**df$**ynose)
dfpoints **<**− setNames(dfpoints, c(“a1”,“a2”,“b1”,“b2”,“y1”,“y2”))
~~~

~~~
*# lengths of the vectors a (tail to neck), b (neck to nose) and y (nose to tail)*
dflength **<**−**data.frame**(**sqrt**(dfpoints**$**a1^2+dfpoints**$**a2^2),
**sqrt**(dfpoints**$**b1^2+dfpoints**$**b2$2), **sqrt**(dfpoints**$**y1^2+dfpoints**$**y2^2))
dflength **<**− setNames(dflength, **c**(“a”,“b”,“y”))
~~~

~~~
#Dot product of a and b
dfab **<**− **data.frame**(dfpoints**$**a1*dfpoints**$**b1+dfpoints**+**a2*dfpoints**\***b2)
dfab **<**− setNames(dfab, c(“ab”))
~~~

~~~
*# Multiplying lengths of a and b*
dfabl **<**− **data.frame** (dflength**$**a*dflength**\***b)
dfabl **<**− setNames ( dfabl, **c** (“abl”))
~~~

~~~
*# Cos of the angle between vectors a and b*
dfanglerad **<**− **data.frame** ( **acos** ( dfab**$**ab / dfabl**$**abl))
dfanglerad <− setNames ( dfanglerad, **c** (“angle_rad”))
~~~

~~~
*# Radians into degrees*
dfangle **<**− data.**frame** ( dfanglerad**$**angle_ rad * (180 / pi))
dfangle **<**− setNames ( dfangle, **c** (“angle”))
~~~

~~~
dfvectors <− **data.frame** (**df**, dfpoints, dflength, dfab, dfabl, dfanglerad, dfangle)
**return** (dfvectors)
}
~~~

~~~
*# Dynamics*
**time**_**diff <**− **function** (**df**) {

# *Time difference between each data point ( t1−t 0, t2−t1, t3−t 2, …)*
**time**_**diff**_value <- **data.frame** (**diff** (**df$time**))
**time**_**diff**_value <- setNames (**time**_**diff**_value, **c** (“diff_time”))
~~~

~~~
# Displacement of the tail marker in y−direction
dytail **<**− **data.frame**(**diff**(**df$**ytail))
dytail **<**− setNames(dytail, **c**(“diff_ytail”))
~~~

~~~
*# Displacement of the tail marker in x−direction*
dxtail **<**− **data.frame**(**diff**(**df$**xtail))
dxtail **<**− setNames(dxtail, **c**(“diff_xtail”))
~~~

~~~
*# Tail velocity in y−direction*
ytail_v **<**− **data.frame**(dytail**$diff**_ytail / **time_diff**_value**$diff_time**)
ytail_v **<**− setNames(ytail_v, **c**(“velocity_ytail”))
~~~

~~~
*# Tail velocity in x−direction*
xtail_v **<**− **data.frame**(dxtail**$diff**_xtail/**time_diff**_value**$diff_time**)
xtail_v **<**− setNames(xtail_v, **c**(“velocity_xtail”))
~~~

~~~
*#Total velocity of the tail marker*
v_tail **<**− **sqrt**((xtail_v**$**velocity_xtail+ytail_v**$**velocity_ytail)^2)
~~~

~~~
*#Displacement of the neck marker in y−direction*
dyneck **<**− **data.frame**(**diff**(**df$**yneck))
dyneck **<**− setNames(dyneck, **c**(“diff_yneck”))
~~~

~~~
*#Displacement of the tail marker in x−direction*
dxneck **<**− **data.frame**(**diff**(**df$**xneck))
dxneck **<**− setNames(dxneck, **c**(”diff_xneck”))
~~~

~~~
*#Neck velocity in y−direction*
yneck_v **<**− **data.frame**(dyneck**$diff**_yneck/**time_diff**_value**$diff_time**)
yneck_v **<**− setNames(yneck_v, **c**(“velocity_yneck”))
~~~

~~~
*#Neck velocity in x−direction*
xneck_v **<**− **data.frame**(dxneck**$diff**_xneck/**time_diff**_value**$diff_time**)
xneck_v **<**− setNames(xneck_v, **c**(“velocity_xneck”))
~~~

~~~
*#Total velocity of the neck marker*
v_neck **<**− **sqrt**((xneck_v**$**velocity_xneck+yneck_v**$**velocity_yneck)^2)
~~~

~~~
*#Displacement of the nose marker in y−direction*
dynose **<**− **data.frame**(**diff**(**df$**ynose))
dynose **<**− setNames(dynose, **c**(“diff_ynose”))
~~~

~~~
*#Displacement of the nose marker in x−direction*
dxnose **<**− **data.frame**(**diff**(**df$**xnose))
dxnose **<**− setNames(dxnose, **c**(“diff_xnose”))
~~~

~~~
*#Nose velocity in y−direction*
ynose_v **<**− **data.frame**(dynose**$diff**_ynose/**time_diff**_value**$diff_time**)
ynose_v **<**− setNames(ynose_v, **c**(“velocity_ynose”))
~~~

~~~
*#Nose velocity in x−direction*
xnose_v **<**− **data.frame**(dxnose**$diff**_xnose/**time_diff**_value**$diff_time**)
xnose_v **<**− setNames(xnose_v, **c**(“velocity_xnose”))
~~~

~~~
*#Total velocity of the nose marker*
v_nose **<**− **sqrt**((xnose_v**$**velocity_xnose+ynose_v**$**velocity_ynose)^2)

**df**_dynamics **<**− **data.frame**(**time_diff**_value, v_tail, v_neck, v_nose, ytail_v, xtail_v, yneck_v, xneck_v, ynose_v, xnose_v)

**return**(**df**_dynamics)
}
~~~

~~~
*#Function to plot the angular change across the time*
**plot_data**_f **<**− **function**(**df**, plotname, **path**) {
*#Function to plot the displacement of the nose in x and y directions*
**plot**_displacement_nose **<**− **function**(**df**,plotname){
*#Displacement*
pn **<**− ggplot() + geom_line(**data** = **df**, aes(x = **time**, y = xnose, group = 1, colour =
“x−direction”)) + geom_line(**data** = **df**, aes(x = **time**, y = ynose, group = 1, color =
“y−direction”)
)
~~~

~~~
*#X & Y labels*
pn **<**− pn+xlab(“Time”) + ylab(“x␣(cm),␣y␣(cm)”) + labs(colour = “Markers”)
*#Title on the middle*
pn **<**− pn + ggtitle(**gsub**(“_”,“␣”,plotname))
pn **<**− pn + theme(**plot.title** = element_**text**(hjust = 0.5))
ggsave(filename=**paste**(**path**, plotname, “_plotnose.png”, sep=“”)
, **plot**=pn, width = 8, height = 6)
}
~~~

~~~
*#Function to plot the angular change of all the rats in the same graph*
**plot**_**all**_angles **<**− **function**(dataframes, **path**) {
p **<**− ggplot() + xlab(“Time␣(ms)”) + ylab(“Angle␣(degree)”)
**for** (iin1:**length**(**file.names**)){
p **<**− p + geom_point(aes_string(x=**get**(**paste**(“df”, **strsplit**(dataframes[i], “.txt”)[1], sep =
“_”))**$time**, y=**get**(**paste**(“df”, **strsplit**(dataframes[i],”.txt”)[1], sep=“_”))**$**angle), size=0.5)
 p **<**− p+ggtitle(“Angular␣change”)
}
ggsave(filename=**paste**(**path**, “plot_all_angles.png”, sep=“”),
**plot**=p, width = 8, height = 6)
}
~~~

~~~
*#To find the set of maximum values of the ytail and corresponding values from the angle
column and calculate its mean value*
**mean**_value_top **<**− **function**(**df**) {
**find**_**max**_ytail **<**− **which**(**df$**ytail == **max**(**df$**ytail))
**all**_**max** <- **df$**angle[**find**_**max**_ytail]
meanvalue <- **mean**(**all**_**max**)
**return**(meanvalue)
}
~~~

~~~
*#Exporting the files in Excel*
savefile.xlsx **<**− **function**(**file**, counter) {
**print**(“Exporting␣vectors…”)
**if** (**file.exists**(**file**)) **file.remove**(**file**)
**for**(i in 1:**length**(counter)) {
**write**.xlsx(**get**(**paste**(“df”, **strsplit**(counter[i], “.txt”)[1], sep = “_”)), **file**,
sheetName=**paste**(**gsub**(“/”,“”,**gsub**(“.txt”,“”, counter[i]))), **append**=TRUE)
}
**print**(**paste**(“Workbook”, (**paste**(“Vectors_”, **basename**(**path**), “.xlsx”, sep=“”)), “has”,
**length**(counter), “worksheets.”))
}
savefile2.xlsx **<**− **function**(**file**) {
**print**(“Exporting␣mean␣values․.”)
**if**(**file.exists**(**file**))**file.remove**(**file**)
**write**.xlsx(**df_max**, **file**, sheetName=**basename**(**path**), **row.names** = FALSE)
}
savefile3.xlsx **<**− **function**(**file**,counter) {
**print**(“Exporting␣dynamics…”)
**if** (**file.exists**(**file**))**file.remove**(**file**)
**for**(i in 1:**length**(counter)) {
**write**.xlsx(**get**(**paste**(“df_dynamics”,**strsplit**(counter[i],“.txt”)[1], sep = “_”)), **file**,
sheetName=**paste**(**gsub**(”/”,“”,**gsub** (“.txt”,“”,counter[i]))), **append**=TRUE)
#addDataFrame(get(paste(“df_acc_”,strsplit(counter[i],“.txt”)[1],sep=“_”)),sheet,row.names=F
ALSE,startRow=1,startColumn=4)
}
**print**(**paste**(”Workbook”, (**paste**(“Dynamics_”,**basename**(**path**), “.xlsx”, sep=“”)), “has”,
**length**(counter), “worksheets.”))
}
run_script **<**− **function**(**path_dir**) {
**path**<**<**−**path_dir
file.names**<**<**− **dir**(**path**, recursive=TRUE, pattern=“.txt”)
*#Run function for each TXT file and name it with df_filename and plot angle vs time*
**for** (i in 1:**length**(**file.names**)) {
**cat**(“processing␣data␣file:”, **file.names**[i], sep=“\n”)
**df**_loop **<**− **read.table**(**paste**(**path**, **file.names**[i], sep=“”))
**df**_loop_**vector <**− process_**data**_f(**df**_loop)
assign(**paste**(“df”, **strsplit**(**file.names**[i], “.txt”)[1], sep = “_”),
**df**_loop_**vector**, envir =
.GlobalEnv)
filename **<**−**paste**(**strsplit**(**file.names**[i], “.txt”)[1], sep = “_”)
**plot**_**data**_f(**df**_loop_**vector**, filename, **path**)
**plot**_displacement_nose(**df**_loop_**vector**, filename)
}
**for** (i in 1:**length**(**file.names**)) {
**cat**(“Dataframes␣for␣Dynamics:”, **file.names**[i], sep=“\n”)
**df**_loop_dynamics **<**− **time_diff**(**get**(**paste**(“df”, **strsplit**(**file.names**[i], “.txt”)[1], sep = “_”)))
assign(**paste**(“df_dynamics”,**strsplit**(**file.names**[i], “.txt”)[1], sep = “_”), **df**_loop_dynamics,
envir = .GlobalEnv)
}
~~~

~~~
*#Finding the maximum, minimum and mean values*
**df_max**<<- **data.frame**()
**for**(i in 1:**length**(**file.names**)){
**cat**(“finding␣Max␣values:”, **file.names**[i],sep=“\n”)
~~~

~~~
*#For the angular change*
**mean**_top **<**− **mean**_value_top(**get**(**paste**(“df”, **strsplit**(**file.names**[i], “.txt”)[1], sep = “_”)))
**mean**_value **<**− **mean**(**get**(**paste**(“df”, **strsplit**(**file.names**[i], “.txt”)[1], sep = “_”))**$**angle)
**max**_value **<**− **max**(**get**(**paste**(“df”, **strsplit**(**file.names**[i], “.txt”)[1], sep = “_”))**$**angle)
**min**_value **<**− **min**(**get**(**paste**(“df”, **strsplit**(**file.names**[i], “.txt”)[1], sep=“_”))**$**angle)
~~~

~~~
*#For the segmental change*
**max**_seg_a **<**− **max**(**get**(**paste**(“df”, **strsplit**(**file.names**[i], “.txt”)[1], sep = “_”))$a)
**max**_seg_b **<**− **max**(**get**(**paste**(“df”, **strsplit**(**file.names**[i], “.txt”)[1], sep = “_”))**$**b)
**max**_seg_y **<**− **max**(**get**(**paste**(“df”, **strsplit**(**file.names**[i], “.txt”)[1], sep = “_”))**$**y)
**min**_seg_a **<**− **min**(**get**(**paste**(“df”, **strsplit**(**file.names**[i], “.txt”)[1], sep = “_”))**$**a)
**min**_seg_b **<**− **min**(**get**(**paste**(“df”, **strsplit**(**file.names**[i], “.txt”)[1], sep = “_”))**$**b)
**min**_seg_y **<**− **min**(**get**(**paste**(“df”, **strsplit**(file.names[i], “.txt”)[1], sep = “_”))**$**y)
**mean**_seg_a **<**− **mean**(**get**(**paste**(“df”, **strsplit**(file.names[i], “.txt”)[1], sep = “_”))**$**a)
**mean**_seg_b **<**− **mean**(**get**(**paste**(“df”, **strsplit**(file.names[i], “.txt”)[1], sep = “_”))**$**b)
**mean**_seg_y **<**− **mean**(**get**(**paste**(“df”, **strsplit**(file.names[i], “.txt”)[1], sep = “_”))**$**y)
~~~

~~~
*#For the velocity change*
**max**_v_tail **<**− **max**(**get**(**paste**(“df_dynamics”, **strsplit**(**file.names**[i], “.txt”)[1], sep =
“_”))**$**v_tail)
**max**_v_neck **<**− **max**(**get**(**paste**(“df_dynamics”, **strsplit**(**file.names**[i], “.txt”)[1], sep =
“_”))**$**v_neck)
**max**_v_nose **<**− **max**(**get**(**paste**(“df_dynamics”, **strsplit**(**file.names**[i], “.txt”)[1], sep =
“_”))**$**v_nose)
**min**_v_tail **<**− min(**get**(**paste**(“df_dynamics”, **strsplit**(**file.names**[i], “.txt”)[1], sep =
“_”))**$**v_tail)
**min**_v_neck **<**− min(**get**(**paste**(“df_dynamics”, **strsplit**(**file.names**[i], “.txt”)[1], sep =
“_”))**$**v_neck)
 **min**_v_nose **<**− min(**get**(**paste**(“df_dynamics”, **strsplit**(**file.names**[i], “.txt”)[1], sep =
“_”))**$**v_nose)
**mean**_v_tail **<**− mean(**get**(**paste**(“df_dynamics”, **strsplit**(**file.names**[i], “.txt”)[1], sep =
“_”))**$**v_tail)
**mean**_v_neck **<**− mean(**get**(**paste**(“df_dynamics”, **strsplit**(**file.names**[i], “.txt”)[1], sep =
“_”))**$**v_neck)
**mean**_v_nose **<**− mean(**get**(**paste**(“df_dynamics”, **strsplit**(**file.names**[i], “.txt”)[1], sep =
“_”))**$**v_nose)
**df_max**[i,1] <**<**− **paste**(“df”, **strsplit**(**file.names**[i], “.txt”)[1], sep = “_”)
**df_max**[i,2] <**<**− **mean**_top
**df_max**[i,3] <**<**− **mean**_value
**df_max**[i,4] <**<**− **max**_value
**df_max**[i,5] <**<**− **min**_value
**df_max**[i,6] <**<**− **max**_seg_a
**df_max**[i,7] <**<**− **max**_seg_b
**df_max**[i,8] <**<**− **max**_seg_y
**df_max**[i,9] <**<**− **min**_seg_a
**df_max**[i,10] <**<**− **min**_seg_b
**df_max**[i,11] <**<**− **min**_seg_y
**df_max**[i,12] <**<**− **mean**_seg_a
**df_max**[i,13] <**<**− **mean**_seg_b
**df_max**[i,14] <**<**− **mean**_seg_y
**df_max**[i,15] <**<**− **max**_v_tail
**df_max**[i,16] <**<**− **max**_v_neck
**df_max**[i,17] <**<**− **max**_v_nose
**df_max**[i,18] <**<**− **min**_v_tail
**df_max**[i,19] <**<**− **min**_v_neck
**df_max**[i,20] <**<**− **min**_v_nose
**df_max**[i,21] <**<**− **mean**_v_tail
**df_max**[i,22] <**<**− **mean**_v_neck
**df_max**[i,23] <**<**− **mean**_v_nose
**df_max**<**<**− setNames(**df_max**, **c**(“Rat”, “mean_top”, “mean”, “max”, “min”, “max_seg_a”,
“max_seg_b”, “max_seg_y”, “min_seg_a”,
“min_seg_b”, “min_seg_y”, “mean_seg_a”, “mean_seg_b”, “mean_seg_y”, “max_velocity_tail”, “max_velocity_neck”,
“max_velocity_nose”, “min_velocity_tail”, “min_velocity_neck”, “min_velocity_nose”,
“mean_velocity_tail”, “mean_velocity_neck”, “mean_velocity_nose”))
}
**plot_all**_angles(**file.names**, path)
savefile.xlsx(**paste**(**path**, “Vectors_”,**basename**(**path**), “.xlsx”, sep=“”), **file.names**)
savefile2.xlsx(**paste**(**path**, “Statistics_”, **basename**(**path**), “.xlsx”, sep = “”))
savefile3.xlsx(paste(path, “Dynamics_”, **basename**(**path**), “.xlsx”, sep=“”), **file.names**)
}
~~~

~~~
*#Always add / at the end of the path*
full_**path** = “/Users/vanessamartinslopes/Documents/Master_thesis/E0510_Vanessa_IDPN_Chronic_in_ra
t/Coordinates/”
*#Read all TXT files on directory*
**list**_weeks **<**− **list**.dirs(full_**path**, full.**names** = FALSE, recursive = FALSE)
**for** (l in 1:**length**(**list**_weeks)){
run_script(**paste**(full_**path**, **list**_weeks[l], “/”, sep=“”))
}
~~~

## REFERENCES

Allum JHJ, Carpenter MG (2013) Postural control and the vestibulospinal system. In: Bronstein AM (ed) Oxford textbook of vertigo and imbalance. Oxford University Press, Oxford, pp 35–48

Boadas-Vaello P, Riera J, Llorens J (2005) Behavioral and pathological effects in the rat define two groups of neurotoxic nitriles. Toxicol Sci 88: 456–466.

Beraneck M, Bojados M, Le Séac’h A, Jamon M, Vidal PP (2012) Ontogeny of mouse vestibulo-ocular reflex following genetic or environmental alteration of gravity sensing. PLoS One 7: e40414.

Besnard S, Lopez C, Brandt T, Denise P, Smith PF (2015) Editorial: The vestibular system in cognitive and memory processes in mammalians. Front Integr Neurosci 9: 55.

Brown DJ, Pastras CJ, Curthoys IS (2017) Electrophysiological measurements of peripheral vestibular function-A review of electrovestibulography. Front Syst Neurosci 11: 34.

Chalansonnet M, Carreres-Pons M, Venet T, Thomas A, Merlen L, Seidel C, Cosnier F, Nunge H, Pouyatos B, Llorens J, Campo P (2018) Combined exposure to carbon disulfide and low-frequency noise reversibly affects vestibular function. Neurotoxicology 67: 270–278.

Curthoys IS, Grant JW, Burgess AM, Pastras CJ, Brown DJ, Manzari L. (2018) Otolithic receptor mechanisms for vestibular-evoked myogenic potentials: A review. Front Neurol 9: 366.

Curthoys IS, MacDougall HG, Vidal P-P, de Waele C (2017) Sustained and transient vestibular systems: A physiological basis for interpreting vestibular function. Front Neurol 8: 117.

De Jeu M, De Zeeuw CI (2012) Video-oculography in mice. J Vis Exp 65: e3971.

Dyhrfjeld-Johnsen J, Gaboyard-Niay S, Broussy A, Saleur A, Brugeaud A, Chabbert C (2013) Ondansetron reduces lasting vestibular deficits in a model of severe peripheral excitotoxic injury. J Vestib Res 23: 177–186.

Gaboyard-Niay S, Travo C, Saleur A, Broussy A, Brugeaud A, Chabbert C (2016) Correlation between afferent rearrangements and behavioral deficits after local excitotoxic insult in the mammalian vestibule: a rat model of vertigo symptoms. Dis Model Mech 9: 1181–1192.

Greguske EA, Carreres-Pons M, Cutillas B, Boadas-Vaello P, Llorens J (2019) Calyx junction dismantlement and synaptic uncoupling precede hair cell extrusion in the vestibular sensory epithelium during sub-chronic 3,3’-iminodipropionitrile ototoxicity in the mouse. Arch Toxicol 93: 417–434.

Hunt MA, Miller SW, Nielson HC, Horn KM (1987) Intratympanic injections of sodium arsanilate (atoxil) solution results in postural changes consistent with changes described for labyrinthectomized rats. Behav Neurosci 101: 427–428.

Imai T, Takimoto Y, Takeda N, Uno A, Inohara H, Shimada S (2016) High-speed video-oculography for measuring three-dimensional rotation vectors of eye movements in mice. PLoS One 11: e0152307.

Jones SM, Jones TA (2014) Genetics of peripheral vestibular dysfunction: lessons from mutant mouse strains. J Am Acad Audiol 25: 289–301.

Jones TA, Jones SM, Vijayakumar S, Brugeaud A, Bothwell M, Chabbert C (2011) The adequate stimulus for mammalian linear vestibular evoked potentials (VsEPs). Hear Res 280: 133–140.

King EB, Shepherd RK, Brown DJ, Fallon JB (2017) Gentamicin applied to the oval window suppresses vestibular function in guinea pigs. J Assoc Res Otolaryngol 18: 291–299.

Llorens J, Callejo A, Greguske EA, Maroto AF, Cutillas B, Martins-Lopes V (2018) Physiological assessment of vestibular function and toxicity in humans and animals. Neurotoxicology 66: 204–212.

Llorens J, Demêmes D (1994) Hair cell degeneration resulting from 3,3′-iminodipropionitrile toxicity in the rat vestibular epithelia. Hear Res 76: 78–86.

Llorens J, Demêmes D, Sans A (1993) The behavioral syndrome caused by 3,3′-iminodipropionitrile and related nitriles in the rat is associated with degeneration of the vestibular sensory hair cells. Toxicol Appl Pharmacol 123: 199–210.

Llorens J, Rodríguez-Farré E (1997) Comparison of behavioral, vestibular, and axonal effects of subchronic IDPN in the rat. Neurotoxicol Teratol 19: 117–127.

Lo WC, Chang CM, Liao LJ, Wang CT, Young YH, Chang YL, Cheng PW (2015) Assessment of D-methionine protecting cisplatin-induced otolith toxicity by vestibular-evoked myogenic potential tests, ATPase activities and oxidative state in guinea pigs. Neurotoxicol Teratol 51: 12–20.

Luebke AE, Holt JC, Jordan PM, Wong YS, Caldwell JS, Cullen KE (2014) Loss of α-calcitonin gene-related peptide (αCGRP) reduces the efficacy of the vestibulo-ocular reflex (VOR). J Neurosci 34: 10453–10458.

Luxa N, Salanova M, Schiffl G, Gutsmann M, Besnard S, Denise P, Clarke A, Blottner D (2013) Increased myofiber remodelling and NFATc1-myonuclear translocation in rat postural skeletal muscle after experimental vestibular deafferentation. J Vest Res 23: 187–93.

Ossenkopp KP, Prkacin A, Hargreaves EL (1990) Sodium arsanilate-induced vestibular dysfunction in rats: effects on open-field behavior and spontaneous activity in the automated digiscan monitoring system. Pharmacol Biochem Behav 36: 875–881.

Pasquet MO, Tihy M, Gourgeon A, Pompili MN, Godsil BP, Léna C, Dugué GP (2016) Wireless inertial measurement of head kinematics in freely-moving rats. Sci Rep 6: 35689.

Russell NA, Horii A, Smith PF, Darlington CL, Bilkey DK (2003) Long-term effects of permanent vestibular lesions on hippocampal spatial firing. J Neurosci 23: 6490–6498.

Saldaña-Ruíz S, Boadas-Vaello P, Sedó-Cabezón L, Llorens J (2013) Reduced systemic toxicity and preserved vestibular toxicity following co-treatment with nitriles and CYP2E1 inhibitors: a mouse model for hair cell loss. J Assoc Res Otolaryngol 14: 661–671.

Schlecker C, Praetorius M, Brough DE, Presler RG Jr, Hsu C, Plinkert PK, Staecker H (2011) Selective atonal gene delivery improves balance function in a mouse model of vestibular disease. Gene Ther 18: 884–890.

Sedó-Cabezón L, Jedynak P, Boadas-Vaello P, Llorens J (2015) Transient alteration of the vestibular calyceal junction and synapse in response to chronic ototoxic insult in rats. Dis Model Mech 8: 1323–1337.

Seoane A, Demêmes D, Llorens J (2001a). Pathology of the rat vestibular sensory epithelia during subchronic 3,3’-iminodipropionitrile exposure: hair cells may not be the primary target of toxicity. Acta Neuropathol 102: 339–348.

Seoane A, Demêmes D, Llorens J (2001b). Relationship between insult intensity and mode of hair cell loss in the vestibular system of rats exposed to 3,3’-iminodipropionitrile. J Comp Neurol 439: 385–399.

Serra A, Salame K, Liao K, Leigh RJ (2013) Eye movements, vision, and the vestibulo-ocular reflexes. In: Bronstein AM (ed) Oxford textbook of vertigo and imbalance. Oxford University Press, Oxford, pp 27–33

Soler-Martín C, Diez-Padrisa N, Boadas-Vaello P, Llorens J (2007) Behavioral disturbances and hair cell loss in the inner ear following nitrile exposure in mice, guinea pigs, and frogs. Toxicol Sci 96: 123–132.

Sichel JY, Eliashar R, Plotnick M, Sohmer H, Elidan J (2000) Assessment of vestibular ototoxicity of ear drops by recording of vestibular evoked potentials to acceleration impulses. Am J Otol 21: 192–195.

Vignaux G, Chabbert C, Gaboyard-Niay S, Travo C, Machado ML, Denise P, Comoz F, Hitier M, Landemore G, Philoxène B, Besnard S (2012) Evaluation of the chemical model of vestibular lesions induced by arsanilate in rats. Toxicol Appl Pharmacol 258: 61–71.

Vulovic V, Curthoys IS (2011) Bone conducted vibration activates the vestibulo-ocular reflex in the guinea pig. Brain Res Bull 86: 74–81.

Wallace DG, Hines DJ, Pellis SM, Whishaw IQ (2002) Vestibular information is required for dead reckoning in the rat. J Neurosci 22: 10009–10017.

Yang TH, Liu SH, Young YH (2010) Evaluation of guinea pig model for ocular and cervical vestibular-evoked myogenic potentials for vestibular function test. Laryngoscope 120: 1910–1917.

